# A smart polymer for sequence-selective binding, pulldown, and release of DNA targets

**DOI:** 10.1101/573030

**Authors:** Elisha Krieg, Krishna Gupta, Andreas Dahl, Mathias Lesche, Susanne Boye, Albena Lederer, William M. Shih

## Abstract

Selective isolation of DNA is crucial for applications in biology, bionanotechnology, clinical diagnostics and forensics. We herein report a smart methanol-responsive polymer (MeRPy) that can be programmed to bind and separate single- as well as double-stranded DNA targets. Captured targets are quickly isolated and released back into solution by denaturation (sequence-agnostic) or toehold-mediated strand displacement (sequence-selective). The latter mode allows 99.8% efficient removal of unwanted sequences and 79% recovery of highly pure target sequences. We applied MeRPy for the depletion of *insulin, glucagon, and transthyretin* cDNA from clinical next-generation sequencing (NGS) libraries. This step improved data quality for low-abundance transcripts in expression profiles of pancreatic tissues. Its low cost, scalability, high stability and ease of use make MeRPy suitable for diverse applications in research and clinical laboratories, including enhancement of NGS libraries, extraction of DNA from biological samples, preparative-scale DNA isolations, and sorting of DNA-labeled non-nucleic acid targets.

## Introduction

Materials that enable selective separation of DNA sequences are crucial for many life science applications. Isolation of high-purity DNA is required across a wide range of scales, for analytical and preparative purposes alike.^1–5^ Next-generation sequencing (NGS), for instance, necessitates extraction of DNA from biological samples, enrichment of a subset of target sequences or depletion of interfering library components.^1,6–8^ Despite rapid advancements, high reagent costs and time-consuming sample preparation remain a significant obstacle for the full implementation of NGS in clinical practice.^9^

Several approaches are available to isolate and purify nucleic acids (NA).^1,8,10^ Target sequences can be pulled down from solution via biotinylated probes that are captured by streptavidin-coated solid beads (e.g. *Thermo Scientific* Dynabeads). Enzymatic approaches can be used to selectively amplify desired NA species or digest undesired ones (e.g. *New England Biolabs* NEBNext rRNA Depletion Kit, or *Evrogen* Duplex-Specific Nuclease^11^). Multiple research groups have recently reported innovative solutions that complement and improve on existing technologies.^2–4,7^

Despite the diverse variety of approaches, current methods have critical shortcomings that require trade-offs between material cost, ease of use, versatility, and performance. Commercial kits are expensive,^9,10^ due to high costs of recombinant proteins and enzymes. Solid beads are susceptible to non-specific interfacial adsorption of macromolecules, leakage of surface-attached streptavidin, degradation in presence of reducing agents and chelators, or incomplete release of target molecules. Most assays have limited stability, shelf life, and a narrow range of compatible experimental conditions. It is therefore crucial to engineer less expensive and more robust materials that allow fast and efficient binding, manipulation and release of selectable target sequences.

To address this challenge, we have developed an oligonucleotide-grafted methanol-responsive polymer (*MeRPy*). MeRPy’s development was inspired by SNAPCAR, a recently reported method for scalable production of kilobase-long single-stranded DNA (ssDNA).^3^ MeRPy acts as a ready-to-use macroligand^12^ for affinity precipitation. We show that the polymer can bind one or multiple DNA targets, isolate, and release selected targets back into the medium. MeRPy pulldown is directly applicable to unlabeled DNA, requires only the most basic laboratory techniques and can be completed within a few minutes of experimental time.

## Results and Discussion

MeRPy is a poly(acrylamide-*co*-acrylic acid)-*graft*-oligo(nucleic acid) copolymer (Figure 1c). It can be selectively precipitated by the addition of methanol (Figure 1a,b). The polymer’s carboxylate groups (1 wt%) are crucial to suppress nonspecific binding of free DNA. Grafted oligonucleotides serve as universal *anchor strands* that provide high binding capacity and activity. To define sequences for target capture and release, MeRPy is programmed with *catcher strand* probes that consist of three domains (Figure 1d): (i) an *adapter site*, (ii) a *target binding site, and* (iii) an (optional) *release site*. After targets hybridize to the binding site, the polymer is precipitated. The pellet is then redispersed in water and targets are released, either non-selectively by thermal or basic denaturation, or selectively via toehold-mediated strand displacement (TMSD)^13^.

**Figure 1.**
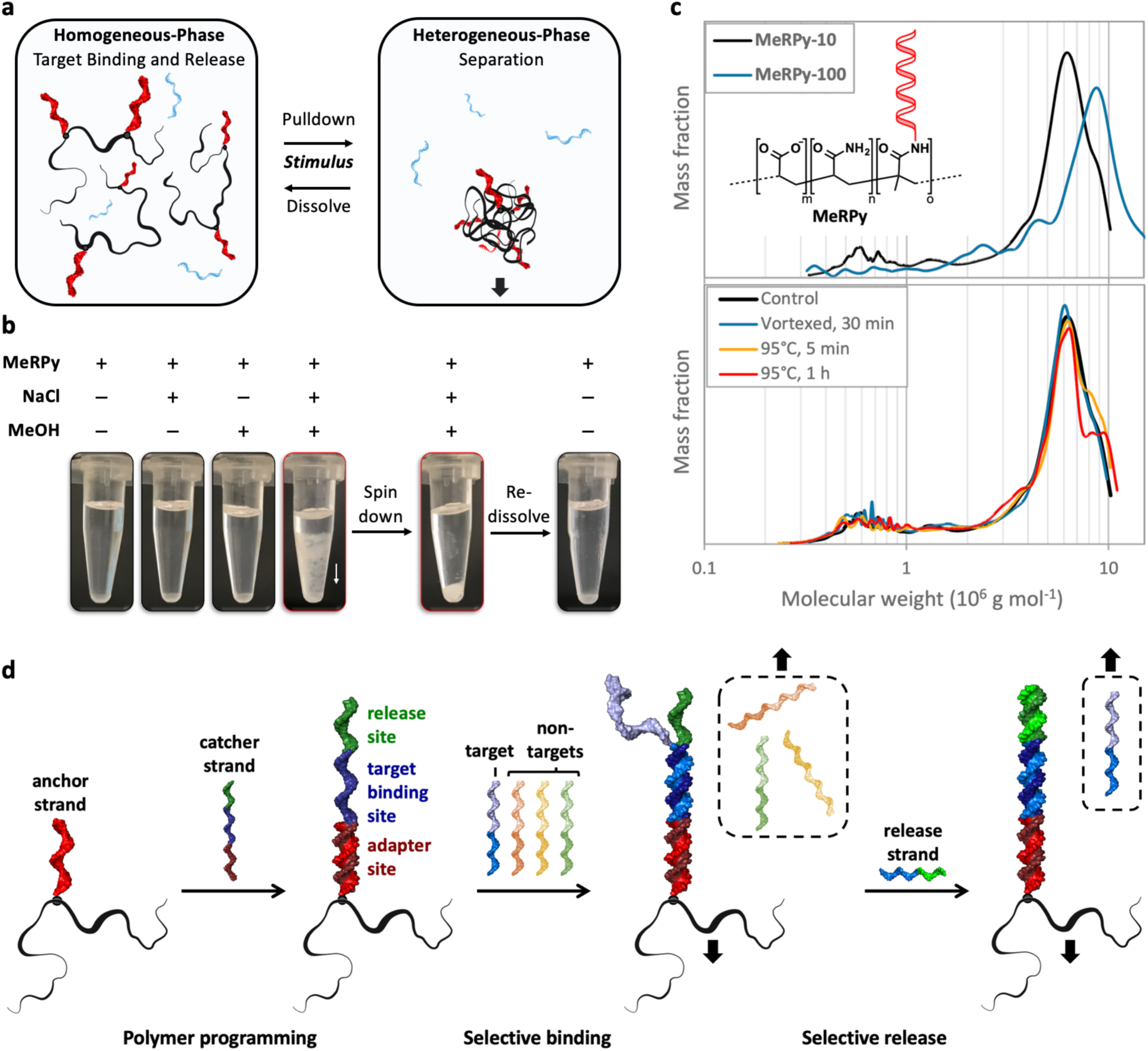
Features of MeRPy. **a)** Due to its methanol-responsiveness, MeRPy combines rapid binding/release in homogeneous phase with facile separation in heterogeneous phase. **b)** MeRPy is soluble in aqueous buffers but precipitates in solutions containing ≥30 mM NaCl when methanol is added. The precipitate can be redissolved in water and triggered to release targets on demand. **c)** Chemical structure and molecular weight distributions. Top: Molecular weight distributions of **MeRPy-10** (m∼740, n∼74,000, o∼10) and **MeRPy-100** (m∼900, n∼90,000, o∼130), obtained by AF4-LS. Bottom: Molecular weight distributions of **MeRPy-10** as synthesized (control), after extensive vortexing and heating to 95°C. **d)** Scheme of MeRPy programming, target binding, and release.

We synthesized two variants of MeRPy (Supplementary Procedure 1): (i) **MeRPy-10** carries ∼10 anchor strands per polymer chain. It can bind up to 2 nmol ssDNA per milligram polymer. (ii) **MeRPy-100** was synthesized for applications that demand increased binding capacity and kinetics. It is endowed with ∼100 anchor strands per chain, providing 20 nmol hybridization sites per milligram polymer. Its binding capacity was found to be 15 nmol per milligram polymer, corresponding to ∼75% of its theoretical limit (Supplementary Figure S1). Both MeRPy variants have significantly higher binding capacities than widely-used magnetic beads, which are limited by molecular crowding at the solid-liquid interface (typically max. 200–500 pmol ssDNA per milligram substrate).^14^

**MeRPy-10** and **MeRPy-100** are soluble in aqueous media. Methanol-induced precipitation requires prior adjustment of the ionic strength, as negative charges of the polymer’s carboxylate groups and anchor strands must be sufficiently shielded by counterions (Figure 1b). **MeRPy-10** requires 30–150 mM NaCl and 1 volume of methanol for complete precipitation. In contrast, **MeRPy-100** requires 100–300 mM NaCl and 1.5 volumes of methanol for this process.

MeRPy’s high molecular weight is crucial for its robust and quantitative responsiveness. Asymmetrical flow-field flow fractionation measurements in combination with static and dynamic light scattering (AF4-LS) indicate that **MeRPy-10** and **MeRPy-100** have weight average molecular weights (M_w_) of 5.73 MDa and 8.47 MDa, respectively. (Figure 1c, Supplementary Figure S3, Supplementary Table S1). The respective dispersities (Ð) are 1.80 and 2.24. MeRPy chains are up to seven times heavier than chains produced in SNAPCAR experiments (M_w_∼1.2 MDa).^3^ The increased molar mass, which improves the efficiency of methanol-induced precipitation, is enabled by its different synthesis approach: in SNAPCAR, target molecules and other reagents can interfere with the *in-situ* radical polymerization; in contrast, MeRPy synthesis is independent of the target capture step, and therefore takes place under highly controlled conditions.

**MeRPy-10** and **MeRPy-100** assume a globular conformation in TE buffer at pH 8.0, as indicated by the scaling exponent (*v*) of 0.38 and 0.32, respectively.^15^ Gyration and hydrodynamic radii support this finding (Supplementary Table S1). The apparent volume (V_app,h_) occupied by individual **MeRPy-10** (V_app,h_ = 2.7×10^−3^ μm^3^) and **MeRPy-100** molecules (V_h_ = 9.2×10^−3^ μm^3^) in solution comprises ≥99.5% water, thus leaving the polymer highly penetrable and anchor strands well accessible for hybridization.

We tested the structural stability of MeRPy under mechanical stress and at high temperature. MeRPy chains remain fully intact when vortexed for 30 minutes or heated to 95°C for 5 minutes (Figure 1c). These exposure times exceed those used in typical pulldown experiments (see below). Heating to 95°C for 1 hour lead to minor decomposition of the upper molecular weight fraction of MeRPy, and partial depurination of anchor strand bases can be expected to occur under these conditions.^16^

To demonstrate that MeRPy is applicable for the separation of DNA and DNA-labeled target molecules, we applied **MeRPy-10** to a mixture of ssDNA-labeled cyanine dyes (T_1_ = Cy5; T_2_ = Cy3) (Figure 2). The polymer was programmed with either of two catcher strands. Fluorescence images show that the targeted dye was selectively pulled down, leaving non-targets in solution. After separating pellet from solution, the release of the captured target into a clean buffer was triggered by the addition of a release strand in presence of 100 mM NaCl. A second MeRPy pulldown then removed the polymer from the released target.

**Figure 2.**
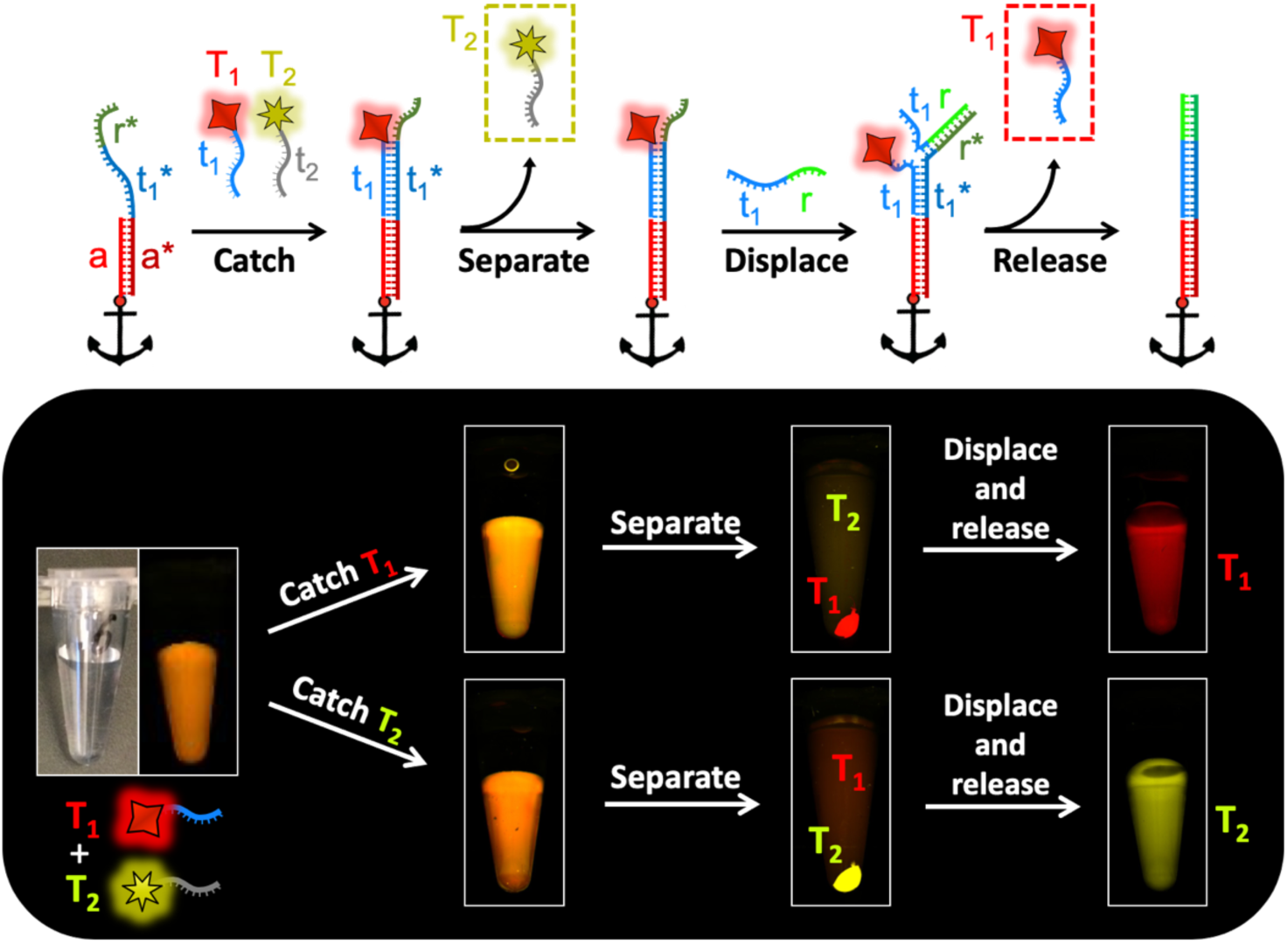
Sequence-specific catch-and-release of ssDNA-labeled target molecules. Photographs of tubes containing **MeRPy-10** and a mixture of two DNA-labeled fluorescent dye molecules. T_1_: *Cy5*; T_2_: *Cy3*. **MeRPy-10** was programmed to bind either T_1_ (upper path) or T_2_ (lower path). The target was then pulled down and isolated by addition of methanol and a short spin-down. After separation, the target was released back into solution by TMSD.

As MeRPy provides high anchor strand concentration, it can be used to capture many targets simultaneously at a fast rate. Figure 3 shows the manipulation of a 10-member ssDNA library (strands designated **A**–**J**) in the length range of 20–190 nt (Supplementary Table S4). Multiple targets were selected by the addition of catcher strand libraries (CSL) of different compositions (Figure 3a). After short annealing of **MeRPy-10** with the target mixture and CSL, the targeted members were depleted from the supernatant with 88±4% efficiency. 98% of non-target strands remained in solution, on average. Nonspecific binding was undetectable for the majority of library components (Figure 3c). Low levels of nonspecific binding were significant only for one library member (strand **H**). After redispersing the pellet in clean buffer solution, the selected library subsets were released via TMSD by addition of either all or only a subset of corresponding release strands. The release efficiency was 90±12%, and the resulting sub-libraries contained the desired strands with a total yield of 79±13%. The recovered target strands were free from non-target contaminations (including strand **H**) within the precision of the measurement (99.8±0.5%). We attribute the high purity to the *dual selection* of target capture and release: target binding first requires the correct binding site sequences (i.e., non-targets stay in the supernatant); TMSD then requires the correct release site sequences (i.e., nonspecifically adsorbed non-targets remain in the pellet).

**Figure 3.**
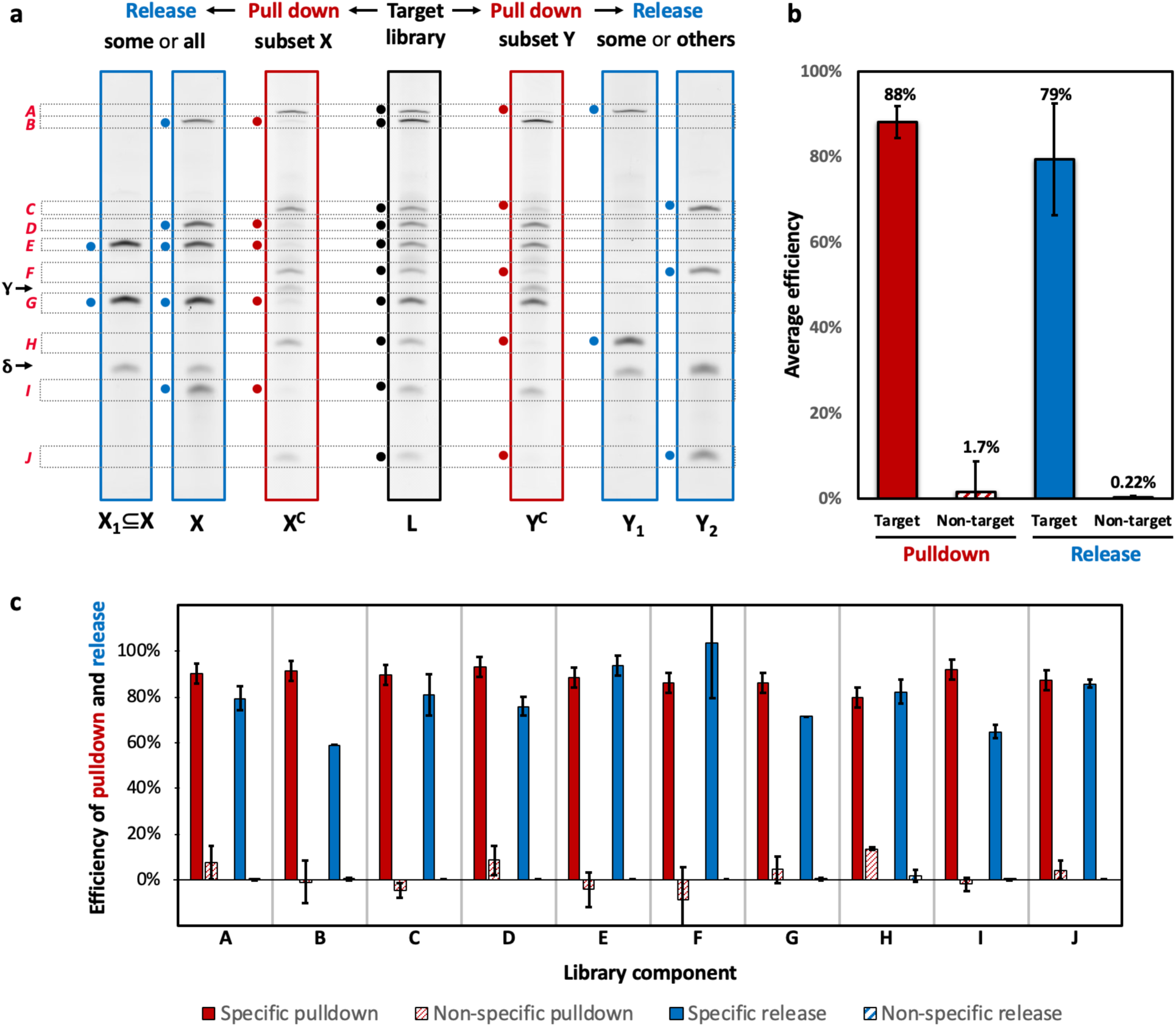
Multiplexed capture and release of selectable targets in a 10-component ssDNA library. **a)** Denaturing polyacrylamide gel electrophoresis (dPAGE) of the library (L) before pulldown (black), after pulldown of selected strands (red), and after release of targeted library subsets (blue). Red and blue circles indicate strands that were targeted by catcher and release strands, respectively. γ = catcher strand band δ = release strand band. **b)** Average binding efficiency and specificity. **c)** Efficiencies and specificities for individual members of the library. Error bars indicate the standard deviation obtained from at least three independent measurements.

In addition to ssDNA, we sought to apply MeRPy to double-stranded DNA (dsDNA) targets. This aim was motivated by the high demand for tools that allow for sequence-selective depletion of complementary DNA (cDNA) from NGS libraries to enhance efficiency and sensitivity in gene expression profiling.^17,7^

Binding dsDNA targets via hybridization is generally challenging, since rapid re-hybridization of target strands after denaturation hampers the sustained attachment of catcher strands. Our approach involved thermal target denaturation, brief annealing with MeRPy and a CSL, followed by rapid MeRPy pulldown. To slow down the rate of sense-antisense rehybridization, the CSL was designed to tile extended regions on the target sequence in an alternating manner (Figure 4a,e). The redundancy of catcher strands ensures that not only full-length targets but also target fragments are depleted.

**Figure 4.**
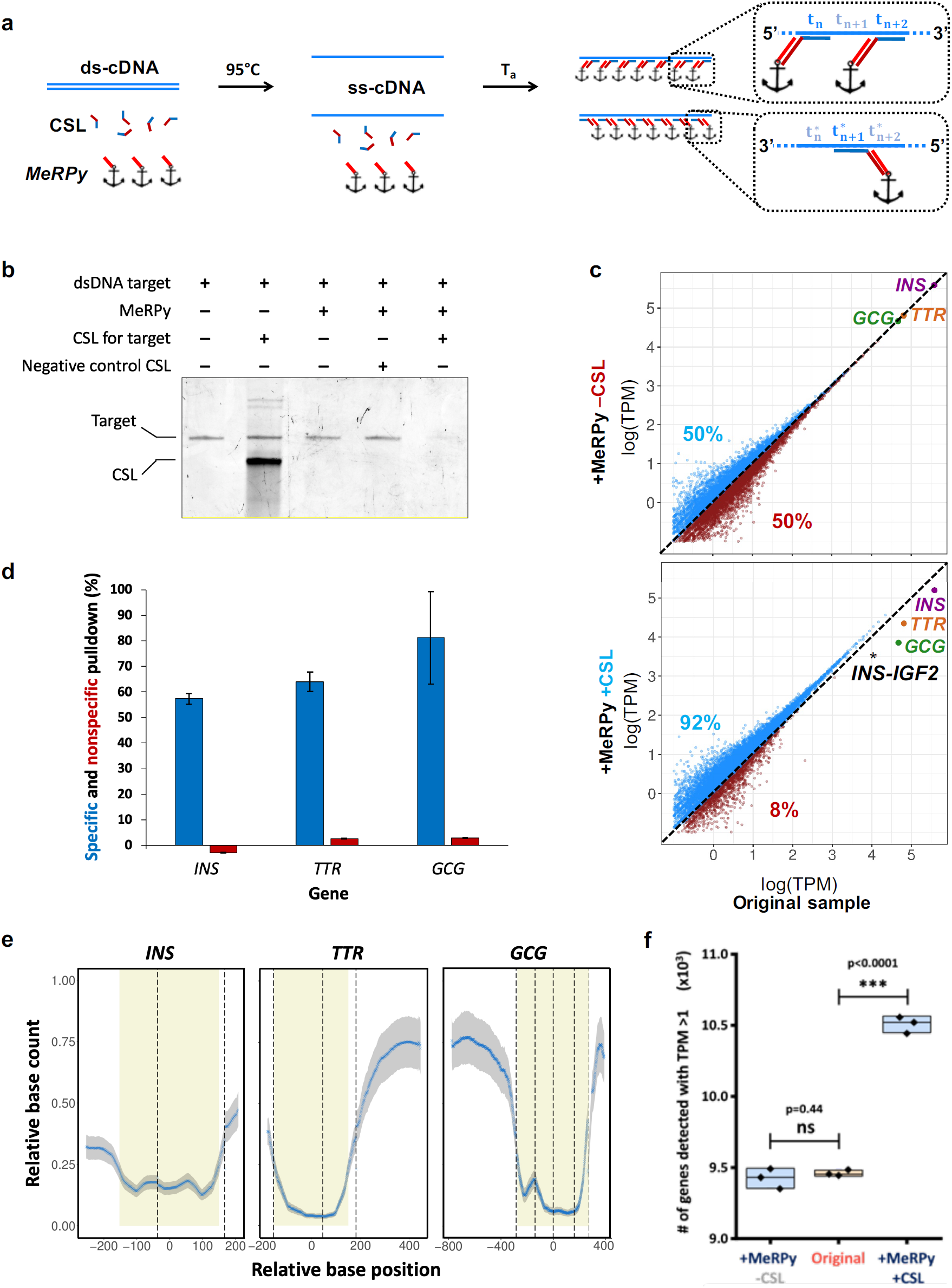
Selective capture of double-stranded DNA (dsDNA) targets. **a)** The target, a catcher strand library (CSL), and MeRPy are mixed and heated to 95°C. Subsequently, the sample is cooled to bind CSL to the target and MeRPy. MeRPy is quickly precipitated to deplete the target from solution. **b)** Denaturing PAGE after pulldown of a dsDNA target (150 bp) with **MeRPy-100**, showing high pulldown efficiency and specificity. **c)** Selective depletion of high-abundance *insulin* (*INS*), *glucagon* (*GCG*), and *transthyretin* (*TTR*) cDNA from a clinical NGS library by MeRPy in presence and absence of an *INS*-, *GCG*-, and *TTR*-targeting CSL. Blue and red data points represent genes that were sequenced with higher and lower number of transcripts per million (TPM), respectively, as compared to the original sample (untreated control). **d)** Pulldown efficiencies for *INS, TTR*, and *GCG*. **e)** Relative base count after pulldown (blue trace) and standard deviation (n=9, grey shade), as a function of base position in the transcripts. CSL-targeted transcript regions are marked in yellow. Relative base positions are relative to the center of the respective targeted region. **f)** Total number of genes detected with >1 TPM in the original sample vs. MeRPy-treated samples (n=3).

Early attempts to capture dsDNA targets with **MeRPy-10** were inefficient. However, **MeRPy-100** allowed up to 10x higher CSL concentrations for increased target binding rates (at the expense of higher material cost; cf. Supplementary Table S2). When using a short annealing protocol (2 min 95°C | 5 min 4°C), followed by immediate pulldown, target binding became favored over sense-antisense rehybridization. Under these conditions, a depletion efficiency of 89.8±2.4% without detectable levels of non-specific binding was achieved for a 150-nt dsDNA mock target strand (Figure 4b).

To demonstrate practical application of the method, we used MeRPy for targeted depletion of highly abundant *insulin* (*INS*), *glucagon* (*GCG*), and *transthyretin* (*TTR*) cDNA from clinical NGS libraries that had been generated from patient-derived pancreatic islets.^18^ Owing to their high expression levels, these three genes consume a large fraction of NGS reads (Supplementary Figure S7), thus affecting the sequencing depth for all other transcripts in the library, some of which carry diagnostically relevant information.^19^

MeRPy pulldown depleted the three targeted genes with 60–80% efficiency (Figure 4c,d). Target depletion was highly selective. MeRPy-treatment did not introduce undesired biases or distortions to the expression profile, as supported by the corresponding Spearman and Pearson correlations (Supplementary figure S9). An unintentional depletion was detected for one non-target gene. The outlier was identified as *INS-IGF2* (Figure 4c), a readthrough gene that aligns to the *INS* sequence, and which was therefore captured by the *INS*-selective CSL.

Within the three depleted genes, base positions that were located around the center of CSL-targeted regions of the transcripts (Figure 4e; yellow regions) were most efficiently depleted, with corresponding base count reduction of 85%, 96% and 94% for *INS, TTR*, and *GCG*, respectively. Regions within the same transcripts that were merely indirectly targeted by the CSL (i.e., by being located upstream or downstream to a targeted region), were also depleted. The reduction in relative base count in these regions decayed with increasing distance to the targeted regions. This effect was expected, as cDNA libraries comprise a wide size distribution of fragments (150–700 nt), some of which lack the CSL-targeted sequence.

Overall, 91% of reads uniquely mapped to the human genome, independent of MeRPy treatment. Therefore, the application of MeRPy and CSL did not generate an interfering background for NGS. The depletion of the three genes from the complex sample made available reads for an additional 327,000 transcripts per million (TPM) (Supplementary Figure S7). As a result, the sequencing depth effectively increased for 92% of genes in the library (Figure 4c). Most importantly, a net surplus of >1000 genes with TPM > 1 were detected in MeRPy-treated samples (Figure 4f).

A similar but less pronounced effect was observed when only one gene, *INS*, was depleted from a cDNA library containing ∼10% *INS*. As described before, high pulldown efficiency (∼80%) and high selectivity were achieved (Supplementary Figure S8). The depletion of *INS* alone increased the sequencing depth for 62% of genes in the library, and a net surplus of ∼350 genes with TPM > 1 were sequenced in *INS*-depleted samples (Supplementary Figures S5, S6).

## Conclusions

In summary, MeRPy enables multiplexable sequence-selective enrichment of DNA targets and depletion of undesired sequences from complex mixtures. Ready-to-use MeRPy solutions contain up to 100 μM anchor strands, thus offering exceptionally high binding capacity and rate, combined with low non-specific adsorption. MeRPy was synthesized for $0.27–0.31 material costs per nanomole binding capacity (Supplementary Table S2). The low cost is a precondition for large-scale and high-throughput applications. Unlike widely used temperature-responsive poly(N-isopropylacrylamide) (PNIPAM)^20–22^ and other DNA-grafted polymers,^23^ MeRPy’s precipitation is triggered by methanol. Its insensitivity towards temperature is important, as it allows for *in-situ* thermal target denaturation and annealing steps that are crucial for achieving high capture efficiency and specificity. MeRPy pulldown is applicable to single-as well as double-stranded DNA targets. Successive target hybridization and TMSD release provided dual sequence-selectivity for retrieval of high-purity target libraries. Depletion of high-abundance cDNA of the genes *INS, TTR*, and *GCG* from NGS libraries was simple and fast (∼10 minutes experimental time). The assay enabled sequencing of low-abundance transcripts in clinical samples that had not been detected in untreated samples at identical sequencing depth. We anticipate that MeRPy will be a useful tool for enhancing the quality and diagnostic value of transcriptomic signatures,^8^ sorting of DNA-encoded chemical libraries,^24^ purification of components for DNA nanotechnology,^25^ as well as isolation of DNA from crude biological samples. Moreover, MeRPy can be used as a stimulus-responsive “macroprimer” in preparative PCR amplifications.^26^

## Methods

### Solvents and reagents

were purchased from commercial sources and used as received, unless otherwise specified. Water was obtained from a *Milli-Q* system. Methanol (MeOH, ACS reagent grade) was obtained from *Fisher Scientific*. Molecular biology grade acrylamide (AA, cat. #A9099), N,N’-methylenebisacrylamide (BAA, cat. #M7279), sodium acrylate (A, cat. #408220), 19:1 acrylamide/bis-acrylamide (cat. #A2917), and ammonium persulfate (APS, cat. #A3678) were purchased from *Sigma Aldrich*. Ultrapure N,N,N’,N’-tetramethylethylenediamine (TEMED, cat. #15524010) and SYBR™ Gold Nucleic Acid Gel Stain (cat. #S11494) were obtained from *Thermo Scientific*. The SequaGel-UreaGel System was purchased from *National Diagnostics* (cat. #EC-833). Desalted oligonucleotides were purchased from *Integrated DNA Technologies* and used without further purification. DNA ladders were purchased from *New England Biolabs* (cat. #N3232L) and Thermo Scientific (cat. #SM0312 and SM1211). Clinical cDNA libraries with concentrations in the range of 1-2 ng/μL were provided by Prof. Michele Solimena (Paul Langerhans Institute Dresden (PLID) TU Dresden). Nitrogen gas (>99.999%) was used for experiments under inert conditions. Nitrogen gas was either supplied by an in-house gas generator, or obtained from pressurized flasks. In the latter case, it was purified through a Model 1000 oxygen trap from *Sigma-Aldrich* (cat. #Z290246). All reagents containing unreacted acrylamide groups were stored at 4°C or −20°C, and protected from unnecessary exposure to light.

### Polyacrylamide gel electrophoresis (PAGE)

was carried out in 0.5x TBE at 150 V on a Mini-PROTEAN^®^ Tetra system from *Bio-Rad Laboratories, Inc.*, using *Serva BluePower*™ *3000* HPE power supply. Denaturing polyacrylamide gels of different percentages were prepared using the SequaGel-UreaGel System. Native polyacrylamide gels were prepared using a 40% (w/w) Acrylamide/Bis-acrylamide (19:1) stock. Gels were stained with SYBR™ Gold. **Gel scans and fluorescence images** were recorded on Typhoon FLA 5100 and Typhoon FLA 9000 Scanners (*GE Healthcare Life Sciences*). Densitometric quantification of gel bands was performed in Fiji (v. 1.0). **UV/Vis absorbance** data was recorded on Nanodrop 1000 and Nanodrop 2000c Spectrophotometers (*Thermo Scientific*). **Sample annealing** was carried out on a CFX96 Touch™ Real-Time PCR Detection System.

### Asymmetrical flow-field flow fractionation with light scattering detection (AF4-LS)

has become an important technique for gentle and detailed characterization of bio-active systems.^27^ We performed the AF4 studies with an Eclipse Dualtec system (Wyatt Technology Europe, Germany). The separation takes place in a long channel (26.5 cm in length), from 2.1 to 0.6 cm in width and a height of 350 μm. The membranes used as accumulation wall were comprised of regenerated cellulose with a molecular weight cut-off of 10 kDa (Merck Millipore DE). Flows were controlled with an Agilent Technologies 1200 series isocratic pump equipped with vacuum degasser. The detection system consisted of a multi angle laser light scattering detector (DAWN HELEOS II from Wyatt Technology Europe, Germany), operating at a wavelength of 658 nm with included QELS module (at detector 99°) and an absolute refractive index detector (Optilab T-rEX, Wyatt Technology Europe GmbH, Germany), operating at a wavelength of 658 nm. All injections were performed with an autosampler (1200 series, Agilent Technologies Deutschland GmbH). The channel flow rate (F_c_) was maintained at 1.0 mL/min for all AF4 operations. Samples (inject load: ∼50 μg) were injected during the focusing/relaxation step within 5 min. The focus flow (F_f_) was set at 3.0 ml/min. The cross-flow rate (F_x_) during the elution step was optimized by an exponential cross-flow gradient of 3 – 0 mL/min in 30 min. 10 mM TRIS buffer and 1 mM EDTA (pH 8.0) was used as eluent for all measurements. Collecting and processing of detector data were made by the Astra software, version 6.1.7 (Wyatt Technology, USA). The molar mass dependence of elution time was fitted with Berry (1^st^ degree exponential).

The apparent volume (V_app,h_) and apparent density (ρ_app,h_) were calculated by equations (1) and (2):

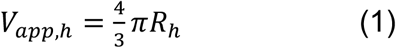

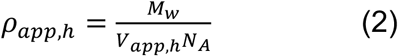

where R_h_ is the hydrodynamic radius, M_W_ is the weight average molecular weight and N_A_ is the Avogadro constant. The scaling exponent was calculated using equation (3)

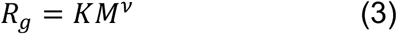

where the scaling exponent (*v*) is a measure for the conformation of the macromolecule. Theoretical values of (*v*) are 0.33 for a hard sphere and 0.5 for a statistical chain in a theta solvent. With increasing macromolecular branching, compact structures with values between 0.5 and 0.33 can be achieved.^28^

### Next-Generation Sequencing (NGS)

#### Initial library preparation and sequencing

Complete cDNA was synthesized from 5 ng total RNA using the SmartScribe reverse transcriptase (Takara Bio) with a universally tailed poly-dT primer and a template switching oligo followed by amplification for 12 cycles with the Advantage 2 DNA Polymerase (Takara Bio). After ultrasonic shearing (Covaris LE220), amplified cDNA samples were subjected to standard Illumina fragment library preparation using the NEBnext Ultra DNA library preparation chemistry (New England Biolabs). In brief, cDNA fragments were end-repaired, A-tailed and ligated to indexed Illumina Truseq adapters. Resulting libraries were PCR-amplified for 15 cycles using universal primers, purified using XP beads (Beckman Coulter) and then quantified with the Fragment Analyzer. Final libraries were subjected to 75-bp-single-end sequencing on the Illumina Nextseq 500 platform.

#### Depletion and sequencing

After depletion of the gene of interest samples were purified using a 1 X volume XP beads (Beckman Coulter), quantified and subsequently subjected to 75-bp-single-end sequencing on the Illumina Nextseq 500 platform.

#### Data analysis

The TPM plots and heatmaps of correlation matrices were generated in R software (v.3.6.1), using ggplot2 and pheatmap packages. The coverage plots were generated by aligning each library and calculating the coverage using bedtools genomecov (v2.27.1) package on Python 3.7 and plotting them on R.

#### Statistics

Statistical significance was tested using an unpaired two-tailed t-test (n = 3, df = 4). The t-test between +MeRPy –CSL and original samples confirmed that the samples had non-significant difference in the number of genes with TPM >1. The t-value and p-value were found to be 0.85 and 0.44 respectively. The t-test between +MeRPy +CSL and original samples resulted in a t-value of 29.0 and p-value <0.0001.

## Acknowledgements

E.K. acknowledges support from the Human Frontier Science Program (LT001077/2015-C) and the Open Topic Postdoc Program of TU Dresden. E.K. would like to thank Prof. Dr. Carsten Werner for support and helpful discussions. W.M.S. acknowledges support from the Wyss Institute at Harvard Core Faculty Award. A.D. acknowledges support from the DFG NGSCC program. We thank Prof. Dr. Michele Solimena for sharing cDNA library samples and the laboratory team of the DRESDEN-concept Genome Center for NGS services.

## Author contributions

E.K. and W.M.S. conceived of the project. E.K. developed MeRPy synthesis, carried out ssDNA pulldown experiments, data analysis and wrote the manuscript. K.G. carried out pulldown experiments with NGS samples, participated in MeRPy synthesis, pulldown protocol optimizations, and data analysis. A.D. and M.L. carried out initial NGS library preparation and sequencing. A.D., M.L. and K.G. carried out sequencing data analysis. S.B. and A.L. performed AF4-LS measurements and data analysis. All authors discussed the results and participated in revising the manuscript draft.

## Additional information

### Competing interests

Provisional patents on this technology have been filed (PCT/US2017/050929 and PCT/US2019/026867).

## Notes

#### Summary of Updates

We collected a comprehensive dataset on cDNA pulldown. In addition to INS, the pulldown experiments now also target the genes TTR and GCG. The multi-target pulldown experiments show significantly improved detection of low-abundance transcripts.

