## Supplementary Information for "A smart polymer for sequence-selective binding, pulldown, and release of DNA targets"

[1]Department of Biological Chemistry and Molecular Pharmacology, Harvard Medical School, Boston, USA; [2]Wyss Institute for Biologically Inspired Engineering at Harvard University, Boston, USA [3]Department of Cancer Biology, Dana-Farber Cancer Institute, Boston, USA; [4]Leibniz-Institut für Polymerforschung Dresden e.V., Germany [5]School of Science, Technische Universität Dresden, Germany; [6]Biotechnology Center (BIOTEC), Technische Universität Dresden, Germany; [7]DRESDEN-concept Genome Center, Center for Molecular and Cellular Bioengineering, Technische Universität Dresden, Germany.

**PROCEDURES** 2

**SUPPLEMENTARY PROCEDURE 1: MERPY SYNTHESIS** 2
1. REAGENTS 2
2. POLYMERIZATION 2
3. PURIFICATION 3
4. STANDARD ANALYSIS 4

**SUPPLEMENTARY PROCEDURE 2: SSDNA PULLDOWN AND RELEASE** 4
1. REAGENTS 4
2. PULLDOWN 5
3. RELEASE 5
4. ANALYSIS 6

**SUPPLEMENTARY PROCEDURE 3: CDNA (DSDNA) PULLDOWN** 6
1. REAGENTS 6
2. PULLDOWN 6

**SUPPLEMENTARY NOTES** 7

**SUPPLEMENTARY FIGURES** 8

**SUPPLEMENTARY TABLES** 12

**SUPPLEMENTARY REFERENCES** 14

### Procedures

#### ***Supplementary procedure 1: MeRPy synthesis***

**1. Reagents**

- Acrylamide-labeled anchor strand (**DNA**), stored at -20°C, protected from light
  - Stock solution 1: 2 mM in TE buffer (~7 kg/mol; ~14 µg/µL)
  - Stock solution 2: 100 µM in TE buffer (~350 g/mol; 700 ng/µL)
- Acrylamide/Sodium acrylate mixture (Label: "**A**"): Acrylamide (**AA**) mixed with Sodium acrylate (**SA**) at 99:1 mass ratio (prepare 40 wt% stock solution in the fume hood; store at 4°C protected from light)
- **TBE** buffer (5x): 500 mM Tris, 500 mM boric acid, 10 mM EDTA, pH 8.2
- **TE** buffer (100x): 1 M Tris, 100 mM EDTA, pH 8.0
- **NaCl**, 5M
- Ultra Low Range DNA ladder (**ULR**)
- Washing solution (**WS**): TE buffer (1x) containing 30 mM NaCl + 1 vol. MeOH
- Tetramethylethylenediamine (**TEMED**, stored at 4°C protected from light)
- Ammonium persulfate (**APS**) powder (stored in sealed tubes at 4°C)
- 2-ml glass vial with septum cap
- Nitrogen gas, >99.999%, passing through a Model 1000 Oxygen Trap
- Formamide-EDTA-BPB loading buffer (**FLB**): 98% Formamide(aq.) containing 5 mM EDTA, 1.7 mM Tris base and 0.02wt% BPB, pH 8.0
- Methanol (**MeOH**)

***2.* Polymerization**

A. Prepare the following solution in a septum-sealed vial (for **MeRPy-10**, see Table 1; for **MeRPy-100**, see Table 2).
   - Mix first TE, TBE, DNA and **A**
   - Degas solution #1 by bubbling with $N_2$ for 20 min
   - In parallel, degas ~2 ml $H_2O$ by bubbling with $N_2$ for 20 min
   - Prepare fresh 10% stock solutions of TEMED and APS:
     - TEMED: 13 µL TEMED + 87 µL degassed $H_2O$ (avoid vortexing; mix gently with pipettor)
     - APS: 100 mg APS + 1 ml degassed $H_2O$ (avoid vortexing; mix gently with pipettor)
   - Add TEMED to sample #1, gently rock vial, then add APS while continuing $N_2$ bubbling

***Table 1.*** *Synthesis of MeRPy with default anchor strand concentration (**MeRPy-10**).*

| ID | TE 1x [µL] | TBE 5x [µL] | DNA 2 mM [µL] | DNA 100 µM [µL] | A 40% [µL] | TEMED 10% [µL] | APS 10wt% [µL] | Total volume [µL] |
|---|---|---|---|---|---|---|---|---|
| **M10** | 498 | 160 | 40 | | 101 | 0.4 | 0.4 | 800 |

| | | | | | | | | |
|---|---|---|---|---|---|---|---|---|
| R | 70.0 | 20 | | 10 | | | | 100 |

*Table 2. Synthesis of MeRPy with 10x higher anchor strand concentration (**MeRPy-100**).*

| ID | TE 1x [µL] | TBE 5x [µL] | DNA 2 mM [µL] | DNA 100 µM [µL] | A 40% [µL] | TEMED 10% [µL] | APS 10wt% [µL] | Total volume [µL] |
|---|---|---|---|---|---|---|---|---|
| **M100** | 138.2 | 160 | 400 | | 101 | 0.4 | 0.4 | 800 |
| **R** | 70.0 | 20 | | 10 | | | | 100 |

B. After adding all reagents, bubble solution gently with $N_2$ for another 30 min
   - The solution should become viscous during that time

C. Retract the $N_2$-supplying needle into the headspace of the vial, tighten screw cap, keep positive $N_2$ pressure in the vial.

D. Incubate sample under $N_2$ for at least 12 h.

E. After incubation, the solution should have turned highly viscous.

F. The vial may be opened to air.

**3. Purification**

A. Using a disposable syringe, take out sample #1 and mix all of it (0.8 mL) with 6.7 mL TE (1x) in a 15 ml Falcon tube.

B. Vortex for at least 10 min. Significantly longer vortexing times may be necessary if the product is highly viscous.

C. Verify that the sample is fully dispersed.

D. Rotate the sample over night in a tube rotator. (***TE_0***)
   - Note: in this protocol **red IDs** indicate that a small fraction of the solution (~50 µL) may be taken out at this point and kept for later analysis.

E. Add to the sample:
   - 25 µL NaCl (5M), if **MeRPy-10** was synthesized
   - 160 µL NaCl (5M), if **MeRPy-100** was synthesized

F. Rapidly add 5.5 mL (0.75 vol.) MeOH and vortex briefly
   - There should be no precipitate. If there's precipitate it should redisperse after further vortexing.

G. Add another 2 ml (0.25 vol.) MeOH and vortex for 20 seconds
   - A white precipitate should form immediately

H. Let sample incubate for 2 min

I. Centrifuge at 100g at r.t. for 5 min and remove supernatant (***S_1***)
   - Note: faster centrifugation will create a pellet that is difficult to redisperse

J. Remove residual supernatant droplets with a pipettor

K. Add $H_2O$ to the pellet until the total sample volume is 7.5 ml

L. Redisperse the pellet by vortexing for 5 min

M. Add 75 µl 100x TE buffer
N. Repeat steps C–L
O. Shake sample for 30 min
P. Verify that the solution is perfectly homogeneous. If it is not, continue shaking.
Q. Split into multiple tubes and store aliquots in the -20°C freezer (**P_2**)

**4. Standard analysis**

A. Only for **MeRPy-100** analysis: dilute samples TE_0, S_1, and P_2 10-fold in TE buffer
B. Prepare samples for polyacrylamide gel electrophoresis (PAGE):

| **Lane** | **1** | **2** | **3** | **4** | **5** | **6** |
|---|---|---|---|---|---|---|
| Sample | 0.1x ULR | R | TE_0 | S_1 | P_2 | 0.1x ULR |
| C (DNA) max. [ng/µL] | 50 | 70 | 75 | 37 | 75 | 50 |
| V (sample) [µL] | 4 | 2 | 2 | 2 | 2 | 4 |
| V (H2O) [µL] | 0 | 2 | 2 | 2 | 2 | 0 |
| V (FLB) [µL] | 16 | 16 | 16 | 16 | 16 | 16 |
| V (loaded) [µL] | 10 | 10 | 10 | 10 | 10 | 10 |

C. Run PAGE:
- 20% Urea-PAGE
- 0.5x TBE running buffer
- Heat samples to 95°C for 20 seconds, then cool down to 4°C prior to loading
- 150V, 45 min
- Stain with SYBR Gold

D. Quantify DNA in samples R, TE_0, S_1, P_2 to determine (i) efficiency of DNA capture, (ii) efficiency of the washing step for removal of free DNA, (iii) final concentration of polymer-attached anchor strands.

### *Supplementary procedure 2: ssDNA pulldown and release*

**1. Reagents**

- Methanol (**MeOH**)
- 10xTE buffer: 100 mM Tris, 10 mM EDTA, pH 8.0
- 1xTE buffer: 10 mM Tris, 1 mM EDTA, pH 8.0
- **TBE** buffer (5x): 500 mM Tris, 500 mM boric acid, 10 mM EDTA, pH 8.2
- **MeRPy-10** (0.5 wt% in TE, ~10 µM max. anchor strand concentration), stored at -20°C
- Target strand library (**TSL**): e.g, target strands #1–10 in Supplementary Table S4 (100 ng/µL each, corresponding to 1.7–16.4 µM; 63.4 µM total oligo concentration).

- Catcher strand library (**CSL**): e.g., a subset of catcher strands #1–10 in Supplementary Table S4 (100 ng/μL each, corresponding to 1.7–16.4 μM)
- Release strand library (**RSL**): e.g., mixtures of release strands #1–10 in Supplementary Table S4 (100 ng/μL each, corresponding to 1.7–16.4 μM)
- 3 M **NaCl**
- Washing solution (**WS**): TE buffer (1x) containing 30 mM NaCl + 1 vol. MeOH
- Formamide-EDTA-BPB loading buffer (**FLB**): 98% Formamide(aq.) containing 5 mM EDTA, 1.7 mM Tris base and 0.02wt% BPB, pH 8.0

**2. Pulldown**

A) To pull down specific targets from a **TSL**, select the appropriate **CSL** and mix together:

| | |
|---|---|
| **MeRPy-10** [μL] | 100 |
| **TSL** [μL] | 3.55 |
| **CSL** [μL] | 7.10 |
| 3M NaCl [μL] | 3 |
| Total volume [μL] | 113.65 |

B) Vortex thoroughly, then anneal:
- 50 °C for 2 min
- Cool to 20°C with a rate of -1.5°C/min

C) Dilute sample with 114 μL 1xTE and vortex shortly

D) Add 228 μL MeOH and quickly mix sample by pipetting up and down
- There should be a precipitate

E) Centrifuge at 100 g for 5 min
- Higher centrifugal force would make the pellet difficult to re-disperse, which would prevent later release of captured targets

F) Retrieve supernatant (**S_1**)

**3. Release**

A) Wash pellet with 500 μL **WS** and spin again at 100 g for 5 min

B) Remove supernatant as well as possible, then add 100 μL H2O to the pellet

C) Vortex for 30 min to fully disperse the pellet

D) Add 10 μL 10xTE

E) Add 3.5 μL 3M NaCl

F) Add 5.5 μL **RSL**

G) Vortex and anneal:
- 50°C for 2 min
- Cool to 25°C with a rate of -1.5°C/min and hold at 25°C for ≥ 1h

H) Add 120 μL MeOH and quickly mix sample by pipetting up and down
- There should be a precipitate

I) Vortex, spin at 4000g for 2 min, then obtain supernatant (**S_2**)

**4. Analysis**

Analyze **S_1** (target-depleted **TSL**) and **S_2** (released **TSL** subset) by dPAGE

- 10% dPAGE
- Dilute 2 µL of sample with 18 µL **FLB**
  - Heat samples to 95°C for 20 seconds, then cool down to 4°C prior to loading
  - Stain with SYBR Gold
  - 150 V, 75 min

### *Supplementary procedure 3: cDNA (dsDNA) pulldown*

**1. Reagents**

- Methanol (**MeOH**)
- **TE** buffer (100x): 1 M Tris, 100 mM EDTA, pH 8.0
- **TBE** buffer (5x): 500 mM Tris, 500 mM boric acid, 10 mM EDTA, pH 8.2
- **MeRPy-100** (0.5 wt% in TE, ~100 µM max. anchor strand concentration), stored at -20°C
- Catcher strand library (**CSL**): 200 µM total oligo concentration, prepared by pooling together equal volumes of catcher strands (200 µM in TE buffer) (cf. Supplementary Table S5)
- **cDNA** (in general: **dsDNA**) samples: concentration: 10-30 ng/µL; tested size range: 150–700 bp; average size: 350–400 bp
- 2.5M **NaCl**
- Formamide-EDTA-BPB loading buffer (**FLB**): 98% Formamide(aq.) containing 5 mM EDTA, 1.7 mM Tris base and 0.02wt% BPB, pH 8.0

**2. Pulldown**

A. Thaw and briefly vortex an aliquot of **MeRPy-100**

B. Prepare the following sample:

| MeRPy-100 [µL] | cDNA [µL] | NaCl (2.5 M) [µL] | CSL (200 µM) [µL] | Total volume [µL] |
|---|---|---|---|---|
| 2 | 2 | 0.5 | 0.5 | 5 |

C. Briefly anneal the sample:
   - Quick heating to 95 °C, hold for 2 minutes
   - Quick cooling to 4 °C, hold for 5 minutes

D. Spin down

E. Immediately add 7.5 µl of MeOH into each solution and quickly pipet up and down to mix the sample, vortex briefly
   - Note 1: Adding the correct volume of MeOH is crucial. The volatility of MeOH may cause the solvent to drip out of the pipette. This problem is best avoided by pre-wetting the pipette tip by repeatedly aspirating and dispensing the solvent (at least 5 times) before transferring the intended volume. Alternatively, a positive displacement pipette may be used.

- Note 2: Avoid any delay between steps C and E, as it may reduce pulldown efficiency due to re-annealing of complementary cDNA fragments.

F. Let the sample stand for 1 min
G. Centrifuge at 2000g for 1 minute
H. Verify that there's a pellet and obtain 9.5 µL of the supernatant (**SN**).

**3. Analysis**

A. Prepare and load following sample for denaturing PAGE (dPAGE)

| SN [µL] | FLB [µL] | Total volume [µL] | Loading volume [µL] |
|---|---|---|---|
| 1 | 9 | 10 | 9 |

B. Run dPAGE:
- 8% dPAGE
- 0.5x TBE running buffer
- Heat samples to 95°C for 20 seconds, then cool down to 4°C prior to loading
- 150V, 45 min
- Stain with SYBR Gold

### Supplementary Notes

AF4 in combination with light scattering detection is a gentle separation and detection technique especially for very complex systems. The separation takes place in a channel. The application of a flow force field allows a controlled separation with reduced shear forces and interactions. The separation range of sizes is much broader (up to 1 µm) compared to routine techniques like size exclusion chromatography (SEC).[1–4] The calculation of the obtained parameters, molecular weight, radius of gyration and hydrodynamic radius allow for the estimation of the scaling properties and the apparent density of the macromolecules. Thus **MeRPy-10** shows a slightly higher scaling exponent ($\nu$ = 0.38-0.39), corresponding to a less compact conformation than **MeRPy-100** ($\nu$ = 0.32). The density calculations confirm higher density for **MeRPy-100** than for **MeRPy-10** (see Table S1). Furthermore, the scaling exponents are typical for rather globular molecular conformation. At the same time the ratio of $R_g/R_h$ of **MeRPY-10** is typical for coil-like structures, highly permeable by the solvent.

### Supplementary Figures

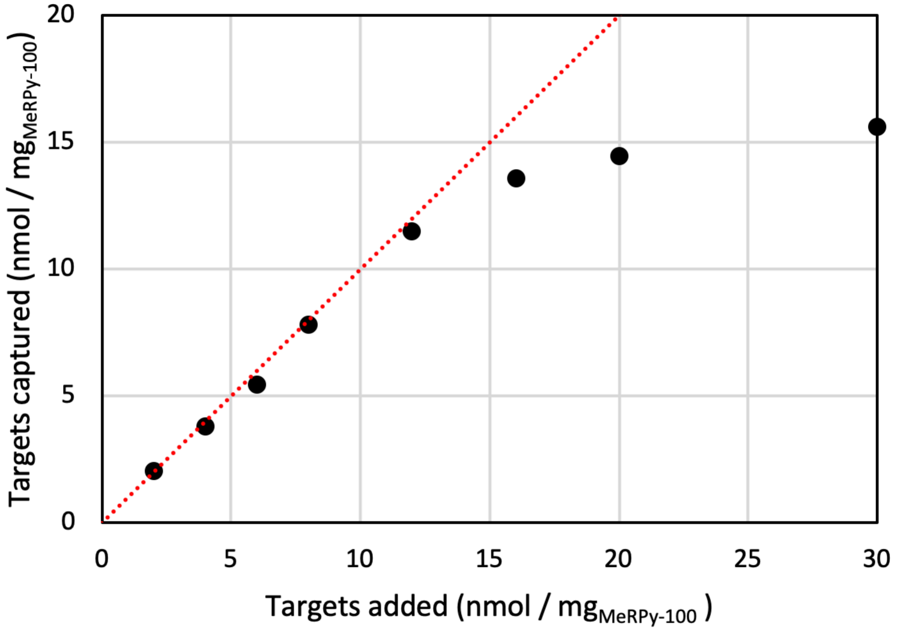


**Supplementary Figure S1.** Experimentally determined binding of catcher strands to **MeRPy-100** (black circles) and theoretical binding limit (red line).

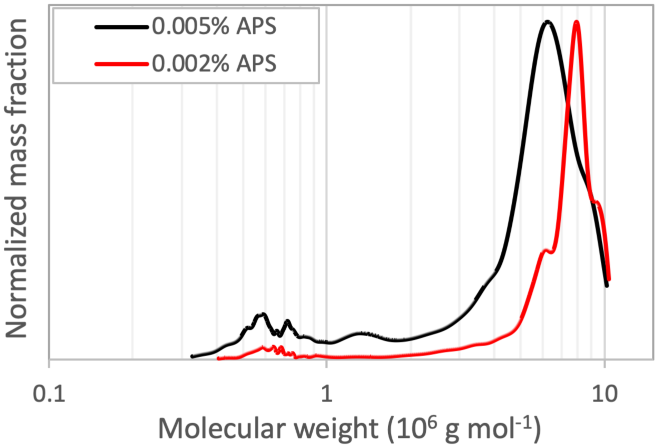


***Supplementary Figure S2.*** *Molecular weight distributions obtained by AF4-LS. Comparison of **MeRPy-10** synthesized with 0.005wt% vs. 0.002wt% of APS.*

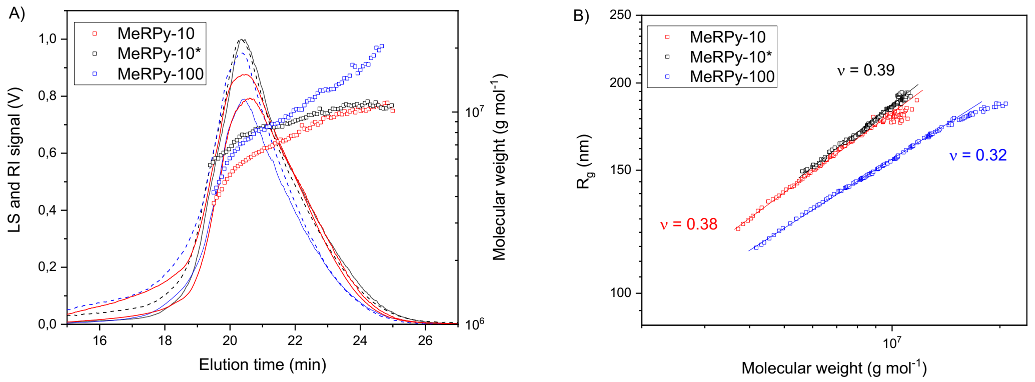


***Supplementary Figure S3.*** *(A) AF4 fractograms: refractive index and light scattering detector signals (solid and dashed lines), molecular weights vs. elution time (squares) of the entire peak region. (B) scaling plots ($R_g$ vs. molecular weight).*

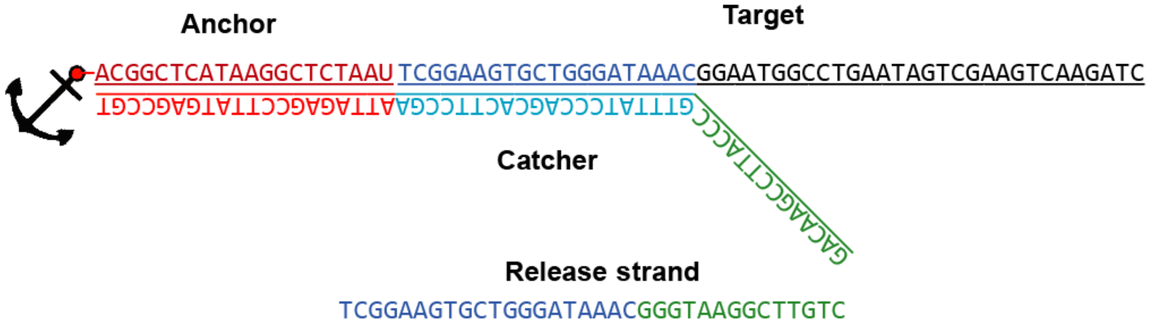


***Supplementary Figure S4.*** *Example of an anchor-catcher-target complex and its corresponding release strand.*

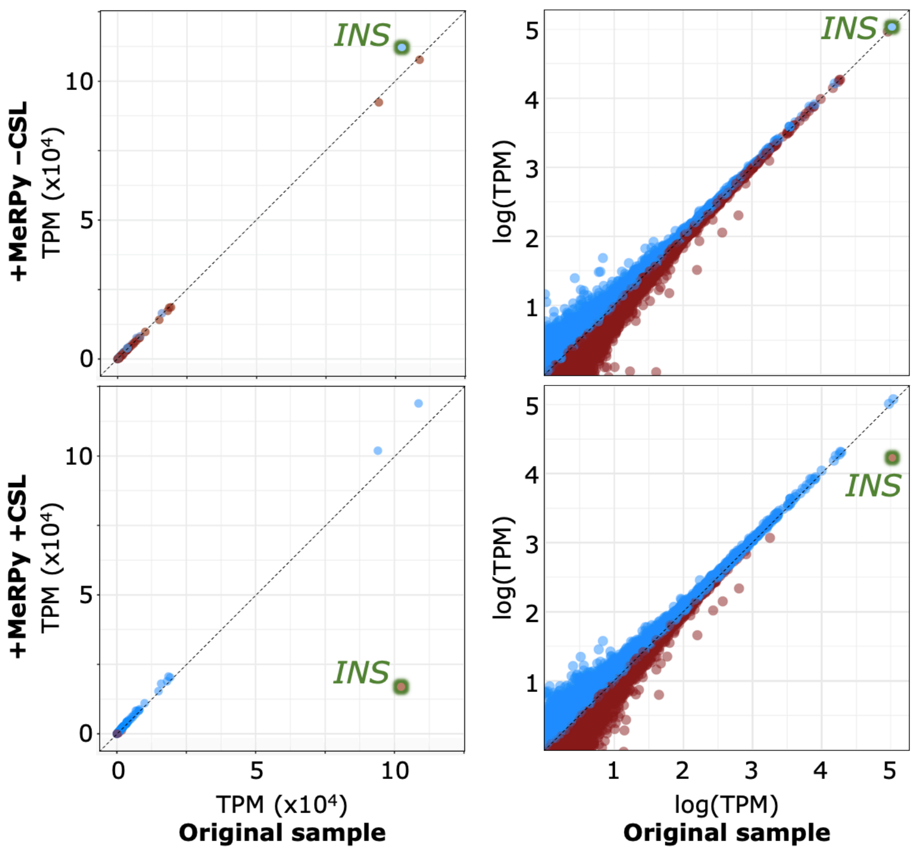


**Supplementary Figure S5.** Selective depletion of high-abundance *insulin* (*INS*) cDNA from a clinical NGS library by MeRPy in presence of an *INS*-specific CSL. Blue and red data points represent genes that were sequenced with higher and lower number of transcripts per million (TPM), respectively, as compared to the original sample.

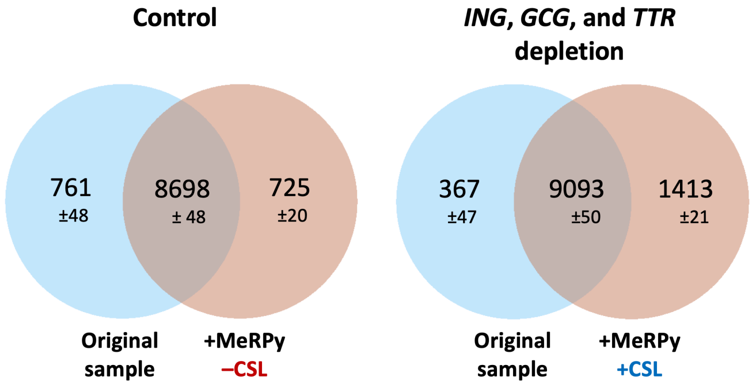


**Supplementary Figure S6.** Venn Diagrams for all genes >1 TPM detection threshold in *INS-*, *GCG-,* and *TTR*-depleted samples (and the –CSL control) vs. the original cDNA library sample.

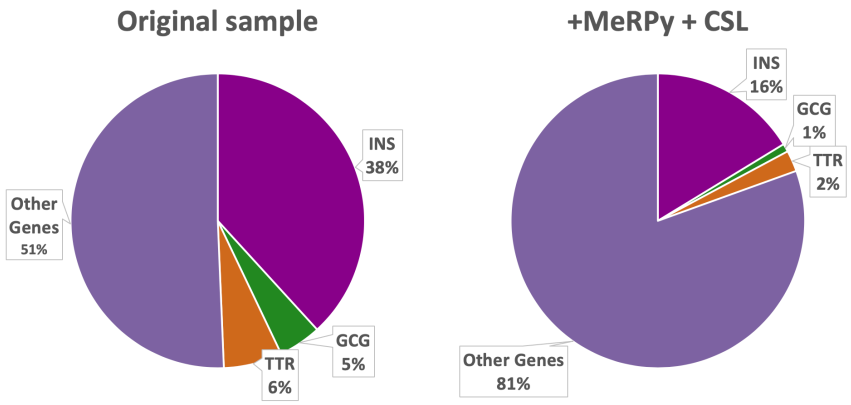


**Supplementary Figure S7.** Fraction of reads consumed by *INS, GCG*, *TTR*, and other genes before (control) and after depletion with MeRPy and a combined *INS*-, *GCG*- and *TTR* -targeting CSL.

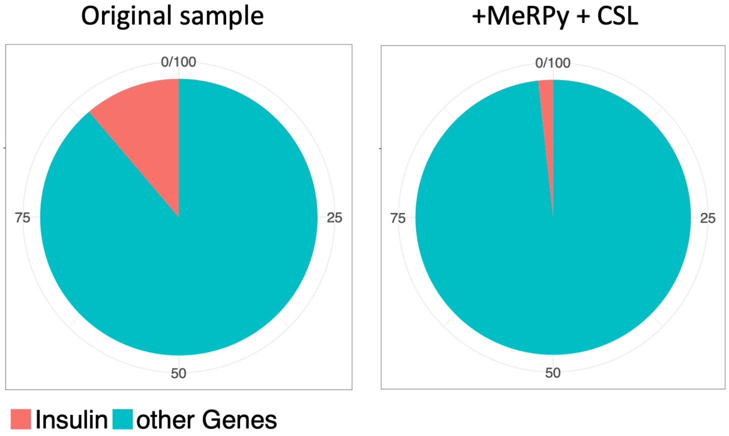


***Supplementary Figure S8.*** Fraction of reads consumed by *INS* and other genes before (control) and after *INS* depletion with MeRPy and an *INS*-specific CSL.

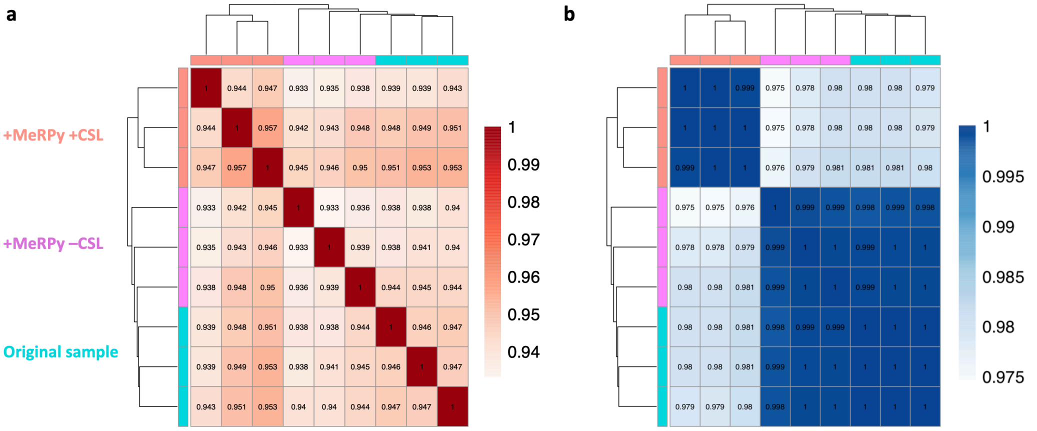


***Supplementary Figure S9.*** **a)** Spearman and **b)** Pearson correlation (all genes, excluding depletion targets*)* for pulldown experiments targeting *INS*, *GCG* and *TTR*.

### Supplementary Tables

***Supplementary Table S1.*** *AF4-LS characterization of **MeRPy-10** and **MeRPy-100**.*

| | $M_W$*1 ($10^6$ g $mol^{-1}$) | Đ*1 ($M_w/M_n$) | $R_g$*2 (nm) | $R_h$*2 (nm) | ν*2 | $R_g/R_h$ *2 | $\rho_{app,h}$ (g/L) | $V_{app,h}$ ($10^{-3}$ $\mu m^3$) |
|---|---|---|---|---|---|---|---|---|
| **MeRPy-10** | 5.73 | 1.80 | 160 | 86 | 0.38 | 1.86 | 4.26 | 2. 7 |
| **MeRPy-10***3 | 7.43 | 1.47 | 173 | 93 | 0.39 | 1.86 | 4.08 | 3.4 |
| **MeRPy-100** | 8.47 | 2.24 | 150 | 130 | 0.32 | 1.15 | 1.27 | 9.2 |

*1 Calculated by dn/dc = 0.160 mL $g^{-1}$, dn/dc for poly acrylamide was determined (0.156 ml $g^{-1}$), and extrapolated for MeRPy with dn/dc for DNA (0.168 ml $g^{-1}$)
*2 Main fraction with high data accuracy
*3 Synthesized with 0.002 wt% (instead of 0.005 wt%) APS and TEMED.
$M_W$ = weight average molecular weight, Đ = dispersity index, $R_g$ = radius of gyration, $R_h$ = hydrodynamic radius, ν= scaling factor (according to Eq. 3), $\rho_{app,h}$ = apparent density calculated by $R_h$, $V_{app,h}$ = apparent volume calculated by $R_h$.

**Supplementary Table S2.** Materials and reagents costs for synthesis and purification of **MeRPy-10** and **MeRPy-100** (40–50 mg scale).

| Materials/reagents | Unit | Unit cost | Cost per prep. MeRPy-10 | Cost per prep. MeRPy-100 |
|---|---|---|---|---|
| Anchor strand | 1 nmol | $0.20 | $16.08 | $160.76 |
| Acrylamide | 1 g | $0.36 | $0.01 | $0.01 |
| Sodium acrylate | 1 g | $2.42 | $0.00 | $0.00 |
| APS | 1 g | $0.42 | $0.00 | $0.00 |
| TEMED | 1 mL | $1.53 | $0.00 | $0.00 |
| $N_2$ gas, purified | 1 $m^3$ | $1.50 | $0.03 | $0.03 |
| 1 ml syringe | 1 | $0.15 | $0.15 | $0.15 |
| 2 ml glass vial | 1 | $0.49 | $0.49 | $0.49 |
| 22G syringe needle | 1 | $0.10 | $0.10 | $0.10 |
| TBE buffer, 5x | 1 mL | $0.06 | $0.01 | $0.01 |
| TE buffer, 100x | 1 mL | $0.45 | $0.09 | $0.09 |
| 1 ml pipette tips | 1 | $0.03 | $0.25 | $0.25 |
| 250 µL pipette tips | 1 | $0.03 | $0.27 | $0.27 |
| 20 µL pipette tips | 1 | $0.03 | $0.55 | $0.55 |
| PCR tube | 1 | $0.05 | $0.30 | $0.30 |
| NaCl, 5M | 1 mL | $0.02 | $0.00 | $0.01 |
| Methanol | 1 L | $6.07 | $0.09 | $0.09 |
| 15 ml Falcon tube | 1 | $0.15 | $0.30 | $0.30 |
| **Total cost** | | | **$18.72** | **$163.41** |
| **Cost per nmol binding capacity** | | | **$0.31** | **$0.27** |

**Supplementary Table S3.** Anchor strand.

| # | Anchor strand | Length [nt] |
|---|---|---|
| 1 | /5Acryd/<u>G</u>ACGGCTCATAAGGCTCTAAX<u>C</u> | 20–22 |

Underlined bases were absent in an early MeRPy synthesis.
X = T or /3deoxyU/

***Supplementary Table S4.*** *Single-stranded DNA target, catcher and release strands.*

| # | Targets | Length [nt] |
|---|---|---|
| 1 | TGTAACATCTGCTGGATCAT | 20 |
| 2 | TGTGCGGACTGGAATGCAAATGAACTGGAT | 30 |
| 3 | GCAAGGATCAAGGTAAAGCTCAGTATAGTCAACGTCAATT | 40 |
| 4 | TCGGAAGTGCTGGGATAAACGGAATGGCCTGAATAGTCGAAGTCAAGATC | 50 |
| 5 | TGCATATCCAGAAGTTCAGTACCCGATCACAGAGTTAGACCATTAGACCATAGCCTTTAC | 60 |
| 6 | TGGACACTGGGATACGAACTTACGAACTTGATCTGATTCTAACTGCCTATAACGAAACTCTATGATAAAA | 70 |
| 7 | TGGTCACGGGTCTCTAAGGTCATAAGTTTCATACGAGGTCAAGGTCAAAGTAGCAATTCCAAGTCAGCGATAAAGTACAA | 80 |
| 8 | CGGTTCATCAAGGTATCAAAGACATCAACTGCCACGTACATAAGCTCGCTCCACTAAGATCCCATTCGTTTAACTAGACC<br>GCTCCAAATC | 90 |
| 9 | GGGATGATGCTGTCGAGCTGGACTCTCACTGTGAACTGAGTTTGGCGTTGCGTTGCGATTATGATCACTCCAATGCATTT<br>TTGCATTGGAGTGATCATAATCGCAACGCAACGCCTTTTGTGCATACAGGTATATCGT | 138 |
| 10 | CGGAGCCGTGCCGCACCGTTCTGTCAGAGTGAGTCAAGCTTTTGGCCCGGCACCCTGCCGCACAATGCCAGCCGTGGTTT<br>ATCAGGCCCAAATAGGTGATATATTTTTATATCACCTATTTGGGCCTGATAAACCACGGCTGGCATTGTGCGGCAGGGTG<br>CCGGGCCTTTGCTCGGAGCTGCCGTCGCCC | 190 |

| # | Catcher strands | Length [nt] |
|---|---|---|
| 1 | CCCTTAGCCACGATATGATCCAGCAGATGTTACAATTAGAGCCTTATGAGCCGT | 54 |
| 2 | CCGTTTTAGATCAGTTTGCATTCCAGTCCGCACAATTAGAGCCTTATGAGCCGT | 54 |
| 3 | ATCTTTGTACACTTAGCTTTACCTTGATCCTTGCATTAGAGCCTTATGAGCCGT | 54 |
| 4 | GACAAGCCTTACCCGTTTATCCCAGCACTTCCGAATTAGAGCCTTATGAGCCGT | 54 |
| 5 | GCTGCTACTTTACGACTGAACTTCTGGATATGCAATTAGAGCCTTATGAGCCGT | 54 |
| 6 | TATCAAGCCGTTAAAGTTCGTATCCCAGTGTCCAATTAGAGCCTTATGAGCCGT | 54 |
| 7 | GTTAGCCAAATGAGACCTTAGAGACCCGTGACCAATTAGAGCCTTATGAGCCGT | 54 |
| 8 | ACCCTGCACTTGACTTTGATACCTTGATGAACCGATTAGAGCCTTATGAGCCGT | 54 |
| 9 | TATTTCAGTTTAGTCAGCTCGACAGCATCATCCCATTAGAGCCTTATGAGCCGT | 54 |
| 10 | TACCTACTGACCCTAACGGTGCGGCACGGCTCCGATTAGAGCCTTATGAGCCGT | 54 |

| # | Release strands | Length [nt] |
|---|---|---|
| 1 | TGTAACATCTGCTGGATCATATCGTGGCTAAGGG | 34 |
| 2 | TGTGCGGACTGGAATGCAAACTGATCTAAAACGG | 34 |
| 3 | GCAAGGATCAAGGTAAAGCTAAGTGTACAAAGAT | 34 |
| 4 | TCGGAAGTGCTGGGATAAACGGGTAAGGCTTGTC | 34 |
| 5 | TGCATATCCAGAAGTTCAGTCGTAAAGTAGCAGC | 34 |
| 6 | TGGACACTGGGATACGAACTTTAACGGCTTGATA | 34 |
| 7 | TGGTCACGGGTCTCTAAGGTCTCATTTGGCTAAC | 34 |
| 8 | CGGTTCATCAAGGTATCAAAGTCAAGTGCAGGGT | 34 |
| 9 | GGGATGATGCTGTCGAGCTGACTAAACTGAAATA | 34 |
| 10 | CGGAGCCGTGCCGCACCGTTAGGGTCAGTAGGTA | 34 |

***Supplementary Table S5.*** *Catcher strand library (**CSL**) for the genes INS, TTR and GCG.*

| # | INS catcher Strands | Length [nt] |
|---|---|---|
| 1 | ATGGCCCTGTGGATGCGCCTCCTGCCCCTGCTGGCGCTGATTAGAGCCTTATGAGCCGTC | 60 |
| 2 | TGAACCAACACCTGTGCGGCTCACACCTGGTGGAAGCTGATTAGAGCCTTATGAGCCGTC | 60 |
| 3 | ACCCAAGACCCGCCGGGAGGCAGAGGACCTGCAGGTGGGATTAGAGCCTTATGAGCCGTC | 60 |
| 4 | CTGCAGCCCTTGGCCCTGGAGGGGTCCCTGCAGAAGCGGATTAGAGCCTTATGAGCCGTC | 60 |
| 5 | CAAAGGCTGCGGCTGGGTCAGGTCCCCAGAGGGCCAGCGATTAGAGCCTTATGAGCCGTC | 60 |
| 6 | GTGTAGAAGAAGCCTCGTTCCCCGCACACTAGGTAGAGGATTAGAGCCTTATGAGCCGTC | 60 |
| 7 | GCTGCCTGCACCAGGGCCCCCGCCCAGCTCCACCTGCCGATTAGAGCCTTATGAGCCGTC | 60 |
| 8 | GGGAGCAGATGCTGGTACAGCATTGTTCCACAATGCCAGATTAGAGCCTTATGAGCCGTC | 60 |
| 9 | GTTCAAGGGCTTTATTCCATCTCTCTCGGTGCAGGAGGGATTAGAGCCTTATGAGCCGTC | 60 |

| # | GCG catcher strands | Length |
|---|---|---|
| 1 | ATGAAAAGCATTTACTTTGTGGCTGGATTATTTGTAATGATTAGAGCCTTATGAGCCGTC | 60 |
| 2 | CAGAGGAGAAATCCAGATCATTCTCAGCTTCCCAGGCAGATTAGAGCCTTATGAGCCGTC | 60 |
| 3 | GCGCCATTCACAGGGCACATTCACCAGTGACTACAGCAGATTAGAGCCTTATGAGCCGTC | 60 |
| 4 | TGGTTGATGAATACCAAGAGGAACAGGAATAACATTGCGATTAGAGCCTTATGAGCCGTC | 60 |
| 5 | CCTTTACCAGTGATGTAAGTTCTTATTTGGAAGGCCAAGATTAGAGCCTTATGAGCCGTC | 60 |
| 6 | AGGAAGGCGAGATTTCCCAGAAGAGGTCGCCATTGTTGGATTAGAGCCTTATGAGCCGTC | 60 |
| 7 | GATGAGATGAACACCATTCTTGATAATCTTGCCGCCAGGATTAGAGCCTTATGAGCCGTC | 60 |
| 8 | TGTCTTGAAGGGAACGTTGCCAGCTGCCTTGTACCAGCGATTAGAGCCTTATGAGCCGTC | 60 |
| 9 | TTGTCCTCGTTCATCTGATCAGGATCACTGAGTGGGTCGATTAGAGCCTTATGAGCCGTC | 60 |
| 10 | CTGCACAAAATCTTGGGCACGCCTGGAGTCCAGATACTGATTAGAGCCTTATGAGCCGTC | 60 |
| 11 | TCCCTTCAGCATGTCTCTCAAATTCATCGTGACGTTTGGATTAGAGCCTTATGAGCCGTC | 60 |
| 12 | CGGCCTTTCACCAGCCAAGCAATGAATTCCTTGGCAGCGATTAGAGCCTTATGAGCCGTC | 60 |
| 13 | AGAGAAAGAACCATCAGCATGTCTGCGGCCAAGTTCTTGATTAGAGCCTTATGAGCCGTC | 60 |
| 14 | CAGTGATTTTGGTCTGAATCAACCAGTTTATAAAGTCCGATTAGAGCCTTATGAGCCGTC | 60 |

| # | TTR catcher strands | Length |
|---|---|---|
| 1 | ATGGCTTCTCATCGTCTGCTCCTCCTCTGCCTTGCTGGGATTAGAGCCTTATGAGCCGTC | 60 |
| 2 | GTGAATCCAAGTGTCCTCTGATGGTCAAAGTTCTAGATGATTAGAGCCTTATGAGCCGTC | 60 |
| 3 | TGTGTTCAGAAAGGCTGCTGATGACACCTGGGAGCCATGATTAGAGCCTTATGAGCCGTC | 60 |
| 4 | GGGCTCACAACTGAGGAGGAATTTGTAGAAGGGATATAGATTAGAGCCTTATGAGCCGTC | 60 |
| 5 | CGGTGCCCGTAGGGCCAGCCTCAGACACAAATACCAGTGATTAGAGCCTTATGAGCCGTC | 60 |
| 6 | TGCACGGCCACATTGATGGCAGGACTGCCTCGGACAGCGATTAGAGCCTTATGAGCCGTC | 60 |
| 7 | ATGCAGCTCTCCAGACTCACTGGTTTTCCCAGAGGCAAGATTAGAGCCTTATGAGCCGTC | 60 |
| 8 | GTGCCTTCCAGTAAGATTTGGTGTCTATTTCCACTTTGGATTAGAGCCTTATGAGCCGTC | 60 |

| # | Dummy adapter sequence | Length |
|---|---|---|
| 1 | TTTTCGCCAAGTAACTCTTTCCGGTTGTTTGCCCGCCAGATTAGAGCCTTATGAGCCGTC | 60 |
